# An EGF-modified PLGA-lanthanide nanoplatform for combined NIR-II cancer imaging and targeted drug delivery

**DOI:** 10.1101/2023.06.18.545497

**Authors:** Yuanyuan He, Zhenfeng Yu, Timo Schomann, Hong Zhang, Christina Eich, Luis J. Cruz

## Abstract

The use of multifunctional nanoplatforms for synergistic therapy and imaging is a promising approach in cancer treatment. In this study, we exploited the imaging properties of lanthanides by encapsulating CaF_2_:Y, Nd along with the chemotherapeutic drug doxorubicin (DOX) into poly (D,L-lactic-co-glycolic acid) (PLGA) nanoparticles (NPs) to prepare a nanoplatform suitable for imaging in the second near-infrared (NIR-II) window and simultaneous anti-cancer therapy. To facilitate the accumulation of CaF_2_:Y, Nd+DOX@PLGA NPs in breast cancer cells, we modified the NPs with EGF. The diameter of the obtained CaF_2_:Y, Nd+DOX@PLGA/PEG/EGF NPs was approximately 150 nm, with a nearly round shape and homogeneous size distribution. In addition, analysis of the drug release behaviour showed that DOX was released more readily and had a longer release time in acidic environments. Accordingly, MTS results indicated that DOX-loaded NPs were significantly cytotoxic. Furthermore, fluorescence microscopy and flow cytometry studies revealed that CaF_2_:Y, Nd+DOX@PLGA/PEG and CaF_2_:Y, Nd+DOX@PLGA/PEG/EGF NPs were gradually taken up by 4T1 breast cancer cells over time, and EGF-coated Nd+DOX@PLGA NPs exhibited increased uptake rates after 72 h. Moreover, we found that EGF increased the solubility of Nd+DOX@PLGA NPs in water by comparing the aqueous solutions of the different NPs formulations. Finally, NIR imaging demonstrated strong fluorescence of PLGA NPs carrying CaF_2_:Y, Nd NPs at 900-1200 nm under 808 nm laser excitation. In conclusion, the developed CaF_2_:Y, Nd+DOX@PLGA/PEG/EGF NPs could be monitored for an extended period of time, and co-encapsulated DOX could be efficiently released to kill breast cancer cells.

## Introduction

Cancer is currently a leading cause of human mortality, mainly due to its high metastasis and recurrence as well as low cure rates[1]. In addition to surgery, most patients require further treatment, such as chemotherapy, radiation, immunotherapy, hormone therapy and targeted therapy[2]. Although drug-assisted therapy is a common cancer treatment, the uncertainty of drug distribution in the body poses potential hazards to the patient. Nanocarriers have emerged as a promising solution to accurately track the drug distribution *in vivo. However*, deep tissue tracking remains a major challenge in current research. Rare-earth-doped nanoparticles (RENPs) exhibit high tissue penetration power and low auto-fluorescence properties in the NIR-II region, making them an effective tool for deep tissue drug tracking[3,4]. Consequently, there is increasing attention and exploration of multifunctional nanoplatforms with tracking, targeting, and therapeutic capabilities.

In recent years, metallic elements have become increasingly prevalent in the preparation of multifunctional nanoplatforms for simultaneous cancer diagnostic and therapy[5]. Lanthanide elements play an important role in this field due to their rich optical, electrical, magnetic and nuclear properties[6]. To date, a large number of lanthanide-based fluorescent materials, such as upconversion nanoparticles (NPs), have been applied for tracking and drug delivery[7]. For example, Kuang *et al. combined* photothermal and radiotherapy nanoplatforms with MRI guidance using the luminescent properties of Gd and the radiosensitizing function of hafnium[8]. Shapoval and his colleagues improved the sensitivity of *in vivo imaging by* doping Nd^3+^ into fluoride contrast agents[9]. A study by Yu *et al. demonstra*ted that Nd^3+^ -doped LuOF nanophosphors (LuOF: Nd^3+^ NPs) displayed excellent NIR-II fluorescence and computed tomography imaging capabilities[10]. Fan *et al. found tha*t modification of RENPs using ligands enhanced their imaging capabilities and showed good tumor-killing capacity when combined with chemotherapeutic agents[11]. However, some limitations, such as the tendency of lanthanides to aggregate and exhibit burst fluorescence, greatly reduce their sensitivity[12]. Fluorides possess unique optical properties (spectral window 190-1100 nm), highly stable structural properties as well as low phonon energies, and are often used as fluorescent host substrates to reduce the bursting of rare earth ions[13]. Recently, we demonstrated that co-doping of calcium fluoride (CaF_2_) with ytterbium (Y^3+^), gadolinium (Gd^3+^), and neodymium (Nd^3+^) broke the quenching threshold of Nd^3+^ and enhanced the NIR-II fluorescence and paramagnetic properties of these multifunctional CaF_2_:Y,Gd,Nd NPs with trimodal imaging capabilities[14].

Doxorubicin (DOX) is a widely used chemotherapeutic agent for the treatment of various types of cancer[15]. As a hydrophilic compound, DOX can inhibit or slow down tumour cell growth *in vivo [16]. Howe*ver, its clinical use is limited by severe cardiotoxicity, drug resistance and damage to healthy tissues, which offsets its survival benefits in cancer patients[17]. To address this issue, PEG-PLGA copolymers have been extensively used for drug delivery due to their biocompatibility, low toxicity, biodegradability and solubility[18-21]. These copolymers can effectively control drug release, improve drug aggregation at the tumor site and reduce side effects[22].

With advances in scientific research, the concept of active targeting has gained significant attention from researchers. NPs can be actively targeted to specific receptors through peripheral ligands[23]. A common ligand-receptor interaction is the binding of epidermal growth factor (EGF) to epidermal growth factor receptor (EGFR), which is overexpressed on cancer cell membranes[24]. EGFR has been shown to promote cancer cell growth, invasion and angiogenesis[25] in numerous cancer cell lines, including lung, colorectal, breast, ovarian, head and neck, prostate, kidney, pancreatic, bladder cancer and glioblastoma[26,27].

To deliver chemotherapeutic drugs while tracking NP distribution *in vivo, we prepar*ed multifunctional NPs by combining CaF_2_:Y,Nd NPs and DOX with PLGA/PEG NPs modified with EGF through a water-in-oil approach. In this platform, Nd^3+^ acts as the main NIR-II imaging centre and Y^3+^ enhances the imaging signal for real-time tracking. The selection of PLGA NPs can locally concentrate CaF_2_:Y,Nd NPs within PLGA NPs, and surface modification with PEG effectively reduces non-specific binding of NPs to blood components, and thus prolongs circulation time *in vivo, and serve*s as a chemical linker to attach targeting motifs to the NP surface. Active targeting and enhanced uptake of CaF_2_:Y, Nd+DOX@PLGA/PEG/EGF NPs were achieved by binding to overexpressed EGF receptors on tumor cell membranes. Here, we analyzed the physicochemical properties of CaF_2_:Y, Nd+DOX@PLGA/PEG/EGF NPs, such as particle size, morphology, and zeta potential, as well as NPs stability, cytotoxicity, *in vitro drug relea*se behavior, cellular uptake and NIR II imaging properties.

## 2 Materials and methods

### 2.1. Materials

Neodymium(III) chloride hexahydrate(NdCl_3_ 6H_2_O, 99.9%), polyvinyl alcohol (PVA, Mw 13,000-23,000, CAS number is 9002-89-5, 87-89% hydrolyzed), chloroform (CHCL_3_, CAS 67-66-3), yttrium (III) chloride heptahydrate (YCl_3_6H_2_O, 99.99%), Calcium chloride dihydrate (CaCl_2_ 2H_2_O, 99.5%), ammonium fluoride (NH_4_F, ≥98.0%), bovine serum albumin and potassium citrate tribasic monohydrate (HOC(COOK)(CH_2_COOK)_2_ H_2_O, ≥99.0%) were obtained from Sigma-Aldrich (Merck KGaA, Darmstadt, Germany). The PLGA polymer (DL-lactide/glycolide molar ratio of 50:50, MW=17,000) was purchased from Carbion PURAC (Amsterdam, the Netherlands). Doxorubicin hydrochloride (DOX, powder, 98.0-102.0 % (HPLC) was acquired from Euroasia Co., Ltd. (Delhi, India). Trypsin, Dulbecco’s modified Eagle’s medium (DMEM) and penicillin-streptomycin (P/S, 10,000 U/mL) were provided by GIBCO (Darmstadt, Germany). Alexa Fluor™ 488 Phalloidin (Cat# A12379), 4’,6-Diamidino-2-Phenylindole (DAPI, Dilactate Cat#D3571), Pierce™ Coomassie (Bradford) Protein Assay Kit (Cat# 23200), and fetal bovine serum (FBS) were purchased from Thermo Fisher Scientific (Waltham, Massachusetts, USA). Agarose was purchased from Bioline (Bioline AgroSciences Ltd., Essex, United Kingdom). CellTiter 96(R) AQueous MTS Reagent Powder were provided by Promega (Madison, Wisconsin, USA). NdCl_3_ (>99%) and YCl_3_ (>99%) were provided by Carlo Erba (Milano, Italia).

### 2.2. Synthesis of CaF_2_:Y, Nd+DOX@PLGA/PEG/EGF NPs

Based on previous studies[28-30], CaF_2_:Y,Nd NPs were prepared by a hydrothermal synthesis process. Briefly, 3.5 mmol stoichiometric CaCl_2_ (>99%, J.T. Baker, Avantor Performance Materials, NJ, United States), YCl_3_ and NdCl_3_ were first added to a 50 mL beaker and dissolved in 7 mL of deionized water. Under vigorous stirring conditions, 20 mL of aqueous potassium citrate (1 M) was added dropwise and after a few minutes, 8.75 mmol of NH_4_F was added and mixed thoroughly. Finally, the clarified mixture was placed in a teflon-lined autoclave, screwed down and baked for 10 h at 180 °C in an oven. After the temperature of the autoclave had been brought to room temperature (RT), the NPs were collected by centrifugation at 2.4 x g for 30 minutes. The samples were washed 4 times with deionized ethanol and water (1:2) to remove any raw materials that did not participate in the reaction or were incompletely reacted, and then freeze-dried for 3 days.

For the preparation of CaF_2_:Y, Nd+DOX@PLGA/PEG/EGF NPs, a double emulsion evaporation technique was used[31,32]. Briefly, 30 mg of DOX and 7.5 mg of CaF_2_:Y, Nd NPs were added to 1 mL of ultrapure deionized water and set aside. Next, 50 mg of PLGA and 10 mg PLGA/PEG-NHS were dissolved in 5 mL of chloroform. After sufficient dissolution, the aqueous solutions of DOX and CaF_2_:Y, Nd NPs were added dropwise to the PLGA chloroform solution. This resulted in the formation of a water-in-oil (W1/O) emulsion, which was emulsified in an ice bath for 65 s using an ultrasonograph (Branson, Danbury, USA). Subsequently, 20 mL of the W1/O emulsion containing 2 % polyvinyl alcohol (PVA, w/v) was added and emulsified by the ultrasonograph to obtain an aqueous W1/O/W2 emulsion. The emulsion was stirred at 4 °C on a magnetic stirrer for 12 h to evaporate the chloroform. The NPs were then collected by high-speed centrifugation at 14,000 x g for 25 min and washed 3 times with deionized water to remove impurities. Next, the NPs were resuspended in 50 μL of EGF solution (1 mg/mL ultrapure deionized water) and stirred at 4 °C overnight. The NPs were collected by high-speed centrifugation (14,000 x g, 25 min) and washed three times to remove the excess EGF. Finally, the obtained NPs were freeze-dried (Martin Christ, Osterode, Germany).

### 2.3. Dynamic light scattering measurements

Zeta potential, particle size distribution and size of the NPs were determined using a Zetasizer Nano ZS90 instrument (Malvern Panalytical, UK). First, 0.5 mg of NPs were dissolved in 1 mL of ultrapure deionized water. The aqueous solution of NPs was then treated in an ultrasonicator for 2 min, and subsequently analyzed using Zetasizer Software (Version 7.13).

### 2.4 Characterization of CaF_2_:Y, Nd NPs

#### 2.4.1 Fourier transform infrared spectroscopy

To analyze the binding properties of CaF_2_:Y,Nd NPs, an IRSpirit FTIR spectrophotometer (Shimadzu, Kyoto, Japan) was used. Briefly, CaF_2_:Y,Nd NP powder was placed into the machine and spectra were recorded in the range 400-4000 cm^-1^, with a resolution of 4 cm^-1^ and 15 scans per spectrum. The data were analyzed after subtraction of the blank control (KBr).

#### 2.4.2 X-ray diffraction

The crystal structure of CaF_2_:Y,Nd NP powder was characterized by X-ray Diffractogram (XRD) analysis. Briefly, the NPs were analyzed using a Panalytical X’pert PRO powder diffractometer (Malvern Panalytical, UK) operating at 40 kV and 40 mA. The analysis was performed at RT with a scanning speed of 6.0 degrees/min over a 2*θ range of 2*0-80 degrees.

#### 2.4.3 Infrared thermal imaging

To examine the effect of photothermal conversion of CaF_2_:Y,Nd NPs under laser irradiation, real-time thermograms and temperature changes after irradiation were recorded using an infrared thermographer (Fluke Ti32, Fluke Corporation, USA)[33-36]. Specifically, 100 μg/mL of CaF_2_:Y,Nd NPs and saline (control) were stored in a 12-well plate. These wells were then irradiated by an 808 nm laser (1.5 W/cm^2^) for 4.5 min and temperature changes were recorded by infrared (IR) thermography during irradiation.

#### 2.4.5 Infrared spectra analysis

The fluorescence of NPs was measured on the Fluorolog^®^ -3 with FluorEssence™ spectrometer (Horiba, Kyoto, Japan). The intrinsic fluorescence was measured by exciting the NP solution at 808 nm (Ex) and obtaining emission (Em) spectra at 900-1200 nm in increments of 0.5 nm.

### 2.6. Drug loading measurements

To measure the drug loading capacity of NPs for the potential antitumor drug DOX, an enzymatic calibrator was used. As described previously[37], 1 mg of NPs was completely dissolved in 0.3 mL of dimethylformamide (DMF) and treated in an ultrasonicator for 30 s to disrupt the structure of the NPs, allowing for complete release of DOX. Next, 0.7 mL of ultrapure water was then added to dilute the solution, and the resulting NP solution was transferred to a 96-well plate. The amount of DOX was analyzed by measuring the absorbance at 480 nm using a microplate reader. The actual concentration of DOX in the NPs was calculated using the following formula:

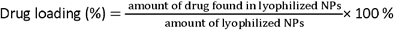

Three independent measurements were performed for this experiment, and data are expressed as mean ± standard deviation (SD).

### 2.7. *In vitro drug relea*se study

The drug release behavior of the NPs was measured using a dynamic dialysis method under different conditions (pH=5.0, 6.5 and 7.4) for 7 days[38]. To accomplish this, an aqueous solution containing 3 mg NPs (with a DOX amount equal to 20 μg) was placed in a dialysis tube (MWCO 3500) and stirred continually in a shaker incubator at 100 rpm and 37 °C. At specific time intervals, 500 μL of the sample was collected and an equal volume of fresh release medium was added. The release of DOX was analyzed at 480 nm using a microplate reader.

### 2.8. Transmission Electron Microscope (TEM)

The morphology and size of DOX-loaded CaF_2_:Y, Nd+DOX@PLGA/PEG/EGF NPs were visualized using transmission electron microscopy (TEM). An aqueous suspension of 1 mg/mL NPs was dropped onto a glow-discharged copper grid surface and the NPs were allowed to settle. The TEM was equipped with a Gatan OneView Camera (Model 1095, Pleasanton, USA) and operated at an accelerating voltage of 120 kV for inspection.

### 2.9. Cell culture

The 4T1 (ATCC No. CRL-2539™) cell line was cultured in DMEM supplemented with 10 % FBS and 1 % P/S and maintained at 37 °C in a 5 % CO_2_ incubator. When the 4T1 cells reached 85-90 % confluence, they were passaged.

### 2.10. *In vitro cytotoxici*ty

The MTS assay was used to assess the cytotoxicity of PLGA/PEG/NPs, PLGA/PEG/EGF NPs, DOX@PLGA/PEG NPs, DOX@PLGA/PEG/EGF NPs, CaF_2_:Y, Nd@PLGA/PEG NPs and CaF_2_:Y, Nd+DOX@PLGA/PEG/EGF NPs. Briefly, 4T1 cells (1 × 10^4^ cells/well) were inoculated into 96-well plates and incubated at 37 °C for 24 h. Subsequently, the cells were treated with different concentrations (31.25, 62.5, 125, 250, 500, 1000 and 2000 μg/mL) of PLGA/PEG NPs, PLGA/PEG/EGF NPs, CaF_2_:Y, Nd @PLGA/PEG NPs, CaF_2_:Y, Nd @PLGA/PEG/EGF NPs, DOX@PLGA/PEG NPs, DOX@PLGA/PEG/EGF NPs, CaF_2_:Y, Nd+DOX@PLGA/PEG NPs and CaF_2_:Y, Nd+DOX@PLGA/PEG/EGF NPs. After 24 and 72 h of incubation, the culture medium was removed and replaced with 100 μL of fresh medium and 20 μL of MTS reagent. The cells were then incubated at 37 °C for 1.5 h. The absorbance of samples was measured at 490 nm using a Molecular Devices SpectraMax M3 Multi-Mode Microplate Reader (Molecular Devices, USA) to assess the cytotoxicity of the NPs.

### 2.11. Bradford Protein Assay

The concentration of EGF protein on the NP surface was measured using the Bradford method[39]. Protein standards with a concentration range of 0.5 to 1000 μg/mL^-1^ were prepared by diluting bovine serum albumin (BSA) in ultrapure water in triplicate according to the manufacturer’s instructions. A 96-well plate was used to mix 50 μL of the solution with 150 μL of Bradford’s reagent, and the mixture was incubation at 37 °C for 30 min. The absorbance of the samples was then measured at 562 nm using a Molecular Devices SpectraMax M3 Multi-Mode Microplate Reader.

### 2.12 NIR-II Fluorescence Imaging

For NIR-II Fluorescence Imaging, 3 mg of PLGA/PEG NPs, PLGA/PEG/EGF NPs, DOX@PLGA/PEG NPs, DOX@PLGA/PEG/EGF NPs, DOX@PLGA/PEG/EGF NPs, CaF_2_:Y, Nd@PLGA/PEG NPs, CaF_2_:Y, Nd@PLGA/PEG/ EGF NPs, CaF_2_:Y, Nd+DOX @PLGA/PEG NPs, and CaF_2_:Y, Nd+DOX @PLGA/PEG/EGF NPs were dissolved in 1.5 mL of ultrapure deionized water. The fluorescence intensity of the aqueous NPs was measured using a laser power density of 100 mW/cm^2^, an excitation wavelength of 808 nm and an exposure time of 5 ms.

### 2.13. Statistical analysis

All data are expressed as mean ± standard deviation (SD) unless stated otherwise. Variance and *t-tests were* used for statistical analysis. A p-value of *p < 0*.05 was considered statistically significant. The symbols *, **, *** and **** denote p-values of *p<0*.*0*5, *p<0*.*0*1, *p<0*.*0*01 and *p<0*.*0*001, respectively. All data analyses were performed using GraphPad Prism 8 software.

## 3 Results and discussion

### 3.1 Characterization of CaF_2_:Y^3+^,Nd^3+^ NPs

Figure 2 depicts representative TEM images of CaF_2_ NPs doped with Y and Nd at different magnifications (Figure 2A). As previously reported[40], our CaF_2_:Y,Nd NPs had uniform morphology and size (Figure 2A), with an average size of 11.8 ± 1.1 nm and a zeta potential of -28.2 ± 0.751, as measured by Malvern Zetasizer Nano (Figure 2B). XRD was used to analyze the crystal structure and dimensions of the NPs, as shown in Figure 2C. An significant peak was observed near the 2*θ value at a* specific angle of 28 degrees, which corresponds to the (111) face of the CaF_2_ phase. In addition, several small peaks were observed near the 2*θ values of* 32 degrees, 67.5 degrees and 76.5 degrees, corresponding to the facets of the (200), (400) and (331) phase, respectively, in line with previous reports[41-43]. The TEM results showed that the crystals ranged between 10 and 15 nm in diameter. Thus, CaF_2_:Y,Nd NPs were highly crystalline, had small grain sizes and exhibited a single-phase cubic fluorite structure.

**Figure 1:**
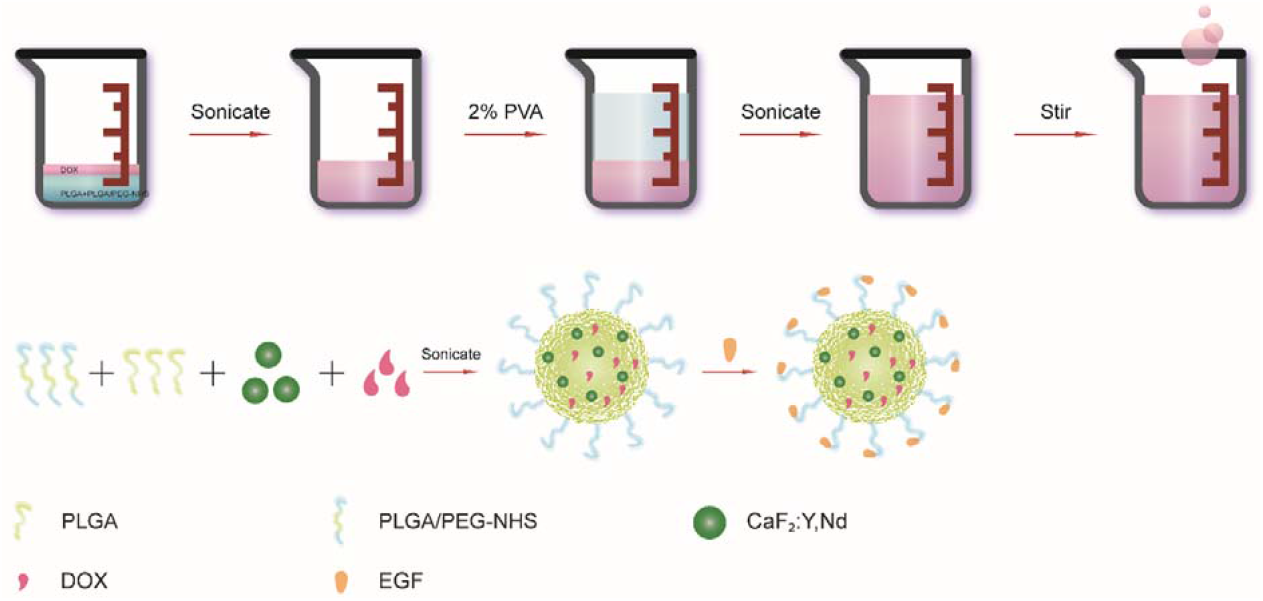
Schematic illustration of the preparation of CaF_2_:Y, Nd+DOX@PLGA/PEG/EGF NPs.

**Figure 2.**
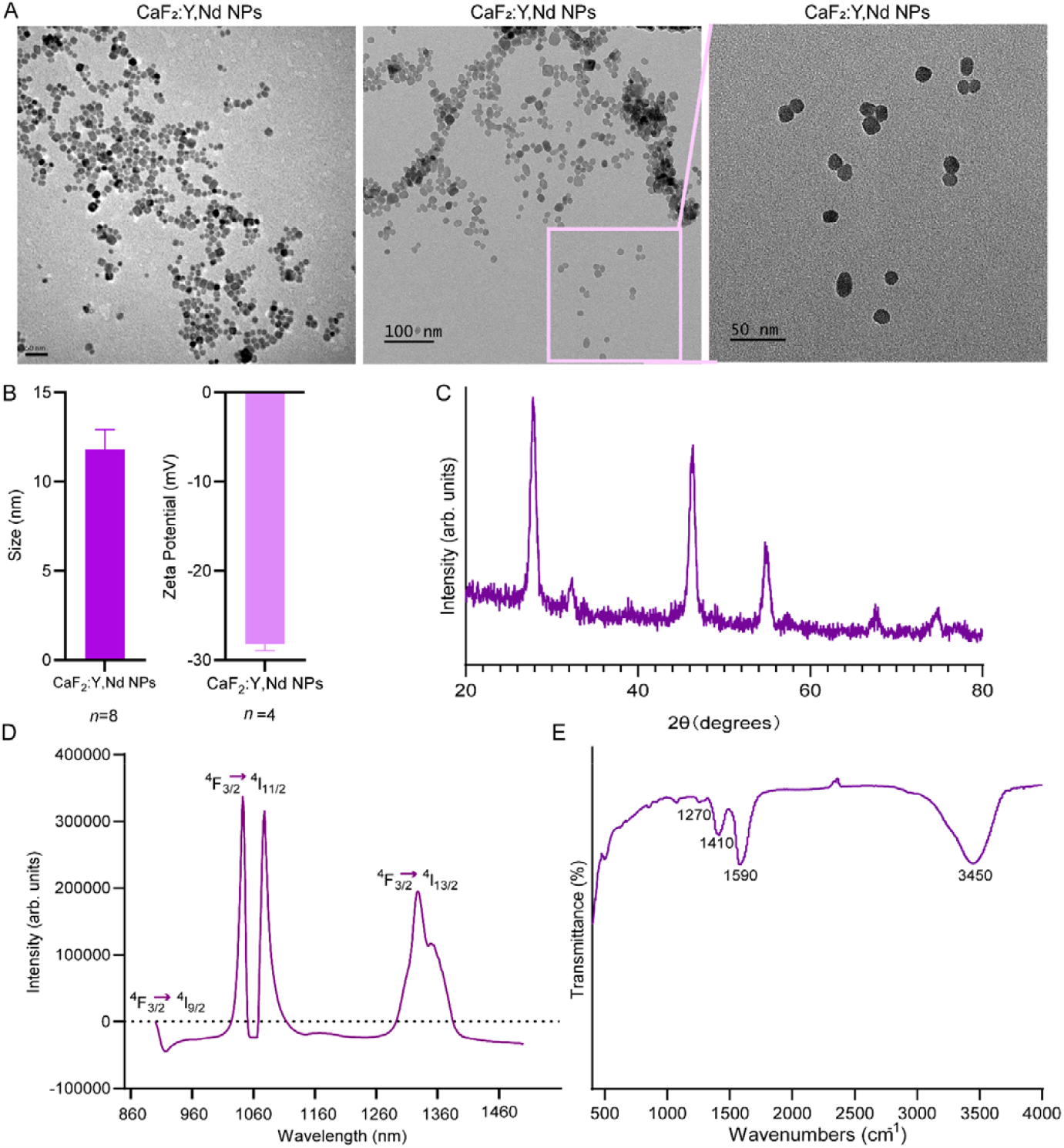
Morphology of CaF_2_:Y,Nd NPs. (A) TEM images of CaF_2_:Y,Nd NPs; (B) Hydrate particle size and zeta potential of CaF_2_:Y,Nd NPs. (C) X-ray diffraction pattern of CaF_2_:Y,Nd NPs. (D) Emission spectra of CaF_2_:Y,Nd NPs (excited by an 808 nm xenon lamp); (E) FT-IR spectra of CaF_2_:Y,Nd NPs.

Next, the CaF_2_:Y,Nd NPs were excited by a NIR diode laser at an excitation wavelength of 808 nm, and strong NIR emission was detected at 1058 nm, 1064 nm and 1330 nm, corresponding to the ^4^F_3/2_ -^4^I_11/2_ and ^4^F_3/2_ -^4^I_13/2_ intra-4f electronic transitions of Nd^3+^, respectively. Since the laser wavelengths fall within the biological first (650-1000 nm) window and the emission wavelengths fall within the biological second (1000-1400 nm) window[28], the NPs can be used for *in vivo imaging of* deep tissues. FT-IR techniques were employed to detect the NPs’ functional groups. As shown in Figure 1E, the adsorption peaks at 1410 and 1590 cm^-1^ corresponded to the stretching vibrations of the carboxyl group in the NPs in the wave number range of 400 to 4000 cm^-1^[44-46]. The absorption peak at 3450 cm^-1^ corresponded to the hydroxyl group stretching vibration in the NPs[47]. In agreement with previous studies, Sisubalan *et al. and Muham*mad *et al. reported* a C-O stretching band at 1410 and 1590 cm^-1^ and an O-H stretching vibration at 3450 cm^-1^, respectively[47,48]. In summary, the properties of our CaF_2_:Y,Nd NPs were consistent with previous results[10,28], demonstrating successful preparation of CaF_2_:Y,Nd NPs, which provided a strong basis for constructing a RENP/PLGA hybrid nanoplatform.

### 3.2 Morphology of CaF_2_:Y, Nd+DOX@PLGA/PEG/EGF NPs

To facilitate the delivery of chemotherapeutic drugs while monitoring the distribution of NPs *in vivo, we prepar*ed CaF_2_:Y, Nd+DOX@PLGA/PEG/EGF NPs using non-toxic degradable PLGA via the compound emulsion method (W1/O/W2). In order to delay drug release, we modified the resulting PLGA-NPs with PEG by a chemical reaction between the carboxyl group (-COOH) of PLGA and the hydroxyl group (-OH) of PEG. To improve the targeting of chemotherapeutic agents and reduce side effects, we further modified the PEG layer of the NPs with EGF via a reaction between the amine-reactive sulfo-NHS ester of PEG and the carboxyl group (-NH_2_) of EGF. To verify the successful encapsulation of CaF_2_:Y,Nd NPs into PLGA NPs, we examined the morphological characteristics of the NPs by TEM. As shown in Figure 3A, the NPs were relatively uniform in size and mostly round. The images of CaF_2_:Y, Nd+DOX@PLGA/PEG/EGF NPs clearly showed the crystal structure of encapsulated CaF_2_:Y,Nd NPs, in line with previous studies that demonstrate the appearance of crystal structures in TEM images as a clear indication of inorganic nanocrystals encapsulated into NPs [49]. The particle size and zeta potential of the NPs were measured using the Zetasizer Nano, which showed that the NPs had a narrow polydispersity index (0.18 ± 0.02) and that their size ranged from 300-600 nm (Figure 3B), with the DLS measurements being similar to the size demonstrated by TEM imaging. In addition, the zeta potential values for PLGA/PEG/EGF NPs, CaF_2_:Y, Nd@PLGA/PEG/EGF NPs, DOX@PLGA/PEG/EGF NPs and CaF_2_:Y, Nd+DOX@PLGA/PEG/EGF NPs were -15.4 ± 0.06 mV, -7.94 ± 0.12 mV, -14.5 ± 0.17mV and -6.67 ± 0.45 mV, respectively (Figure 3C). Our experimental results were consistent with previous research. For example, Lei *et al. reported* a zeta potential of -13.2 ± 2.3 mV for DOX@PLGA NPs [50]. Notably, both NP formulations encapsulating CaF_2_:Y,Nd were less negative than the NPs without lanthanides.

**Figure 3.**
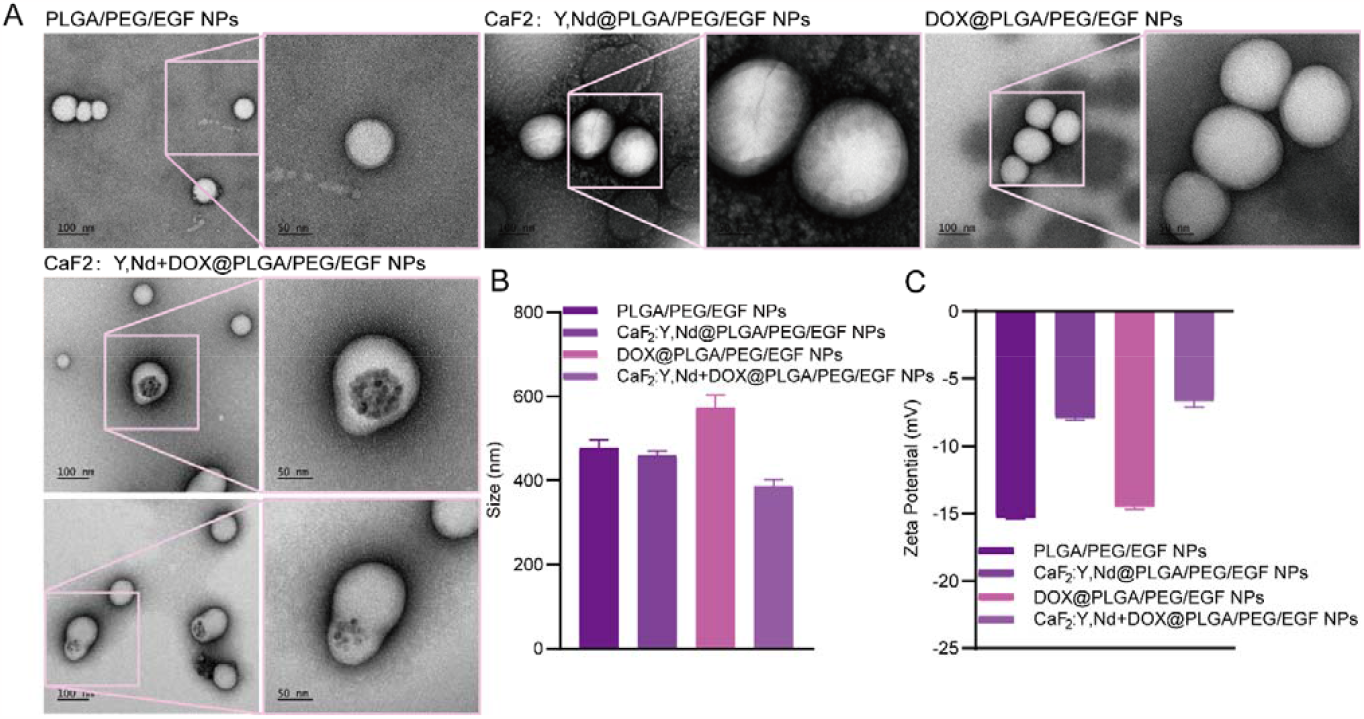
The TEM (A) images and the size distribution (B and C) of NPs in control and experimental groups.

To determine the amount of EGF attached to our CaF_2_:Y Nd+DOX@PLGA/PEG/EGF NPs, we utilized the Bradford Protein Assay [51]. As shown in Supplementary Figure 1, the amount of EGF detected in 1 mg of CaF_2_:Y Nd+DOX@ PLGA/PEG/EGF NPs was 0.93 ± 0.1 μg/mg. In comparison, background levels of 0.086 ± 0.006 μg/mg and 0.048 ± 0.01 μg/mg were detected in CaF_2_:Y Nd+DOX@PLGA/PEG NPs and ultrapure water, respectively. These results validate that EGF was successfully conjugated to the surface of CaF_2_:Y Nd+DOX@PLGA/PEG/EGF NPs.

**Supplementary Figure 1.**
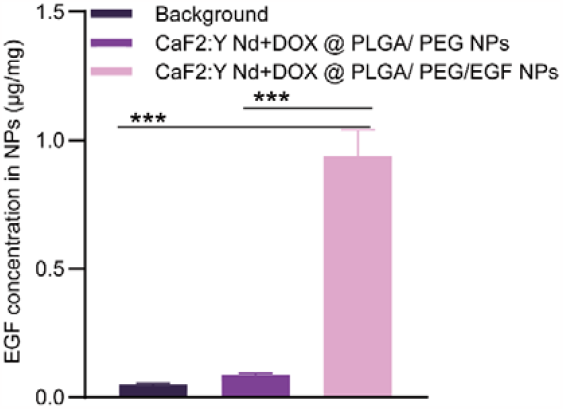
The concentration of EGF protein in NPs was measured using the Bradford protein assay, ***p < 0.001 compared with control group by t-test.

### 3.3. NIR-II spectra Analysis

The NIR fluorescence spectrum of CaF_2_:Y,Nd+DOX@PLGA/PEG/EGF NPs was characterized by the Fluorolog ^®^-3 with FluorEssence™ spectrometer. As shown in Figure 4, PLGA NPs containing CaF_2_:Y,Nd NPs exhibited enhanced fluorescence under 808 nm excitation. In contrast, the fluorescence spectra of PLGA NPs without CaF_2_:Y,Nd NPs showed no peaks. This indicates that the fluorescence performance of CaF_2_:Y,Nd+DOX@PLGA/PEG/EGF NPs was attributed to the doped CaF_2_:Y,Nd NPs. Our previous study confirmed that Y^3+^ and Nd^3+^ could be doped in CaF_2_, forming NPs that emit light stably under 808 nm laser excitation[28]. Several studies have shown that lanthanides can be efficiently loaded into other nanoplatforms while retaining their original fluorescence properties[52-56]. For example, Kang *et al. created m*ultifunctional nanocomposites by loading polyacrylic acid hydrogels with gadolinium vanadate, which acted not only as cellular imaging bioprobes but also facilitated the efficient release of co-loaded DOX in cancer cells[52]. Our data demonstrated that after the addition of DOX, the NPs retained their fluorescence properties, but the fluorescence intensity was slightly reduced (Figure 4). We believe that this reduction was likely due to the co-encapsulation of DOX, which reduced the available space for CaF_2_:Y,Nd, causing a reduction in fluorescence signal in CaF_2_:Y,Nd+DOX@PLGA/PEG/EGF NPs.

**Figure 4.**
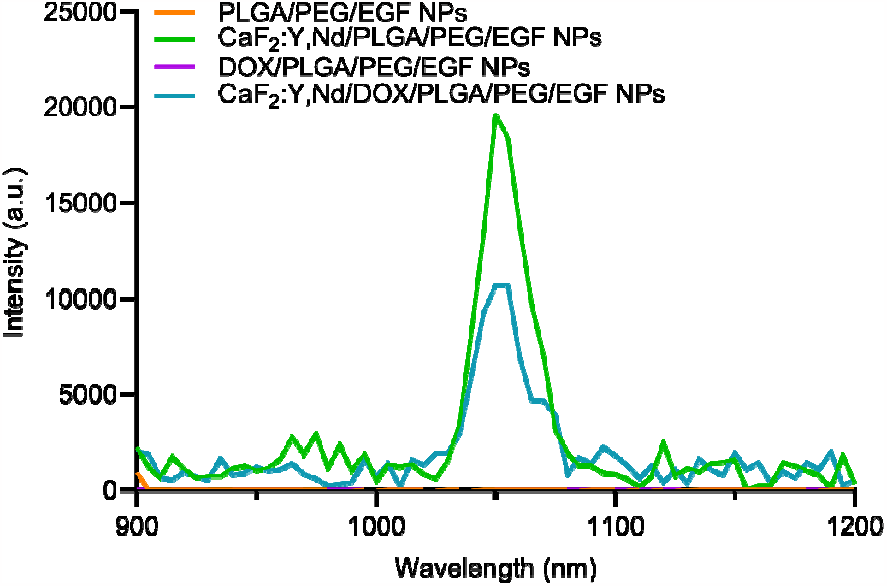
The fluorescence spectra of CaF_2_:Y,Nd+DOX@PLGA/PEG/EGF NPs under 808 nm excitation. Emission spectra of PLGA/PEG/EGF NPs (orange), CaF_2_:Y,Nd@PLGA/PEG/EGF NPs (green), DOX@PLGA/PEG/EGF NPs (purple), and CaF_2_:Y,Nd+DOX@PLGA/PEG/EGF NPs (blue).

### 3.4 Drug release

The therapeutic efficiency of NPs is directly influenced by the efficiency of drug loading. Figure S2 displays the standard curve of DOX at RT at a concentration range of 1.95-500 μg/mL. The loading efficiency of DOX in our NPs was calculated by determining the absorption spectrum at 490 nm of solutions of NPs dissolved in DMF[57-60]. The DOX loading rates were analyzed after adding different amounts of DOX during the NP preparation, as shown in Figure S3. By analyzing the data, we found that the DOX loading rate was proportional to the amount of added DOX. The loading of DOX into CaF_2_:Y,Nd+DOX@PLGA/PEG NPs and CaF_2_:Y,Nd+DOX@PLGA/PEG/EGF NPs was calculated to be 8.4 % and 8.9 %, respectively. This indicates that DOX was efficiently co-encapsulated into PLGA NPs.

Next, the cumulative release of DOX from CaF_2_:Y,Nd+DOX@PLGA/PEG NPs and CaF_2_:Y,Nd+DOX@PLGA/PEG/EGF NPs was investigated over time at different pH values (pH=5.0, 6.5 and 7.4). The pH 5.0 was used to mimic the microenvironment of endosomes and lysosomes[61], pH 6.5 mimicked the weakly acidic microenvironment in tumor cells[62], and pH 7.4 mimicked the physiological environment that NPs encounter in the bloodstream[63]. After 24 h, 63.0 %, 47.5 %, and 35.0 % of drugs were released from CaF_2_:Y,Nd+DOX@PLGA/PEG NPs in PBS at pH 5.0, 6.5, and 7.4, respectively. The drug release was monitored for up to 30 days (720 h) reaching a cumulative DOX release of 65.0 % at pH 5.0, 50 % at pH 6.5, and 36 % at pH 7.4. Under more acidic conditions, the release of DOX was more rapid and the cumulative release was greater, indicating that the pH significantly affected the release of DOX from rapidly in an acidic environment.

Additionally, we observed that the release of DOX from CaF_2_:Y, Nd+DOX@PLGA/PEG NPs was more complete than that from NPs surface-modified with EGF NPs (Figure 5A-D), suggesting that the presence of EGF limited the release of DOX. This finding is line in line with other studies showing that modification with EGF, for example, reduced the curcumin release rate from EGF-modified curcumin/chitosan NPs[65]. Similarly, Wang *et al. also foun*d that cisplatin was slowly released from cisplatin-loaded EGF-modified mPEG-PLGA-PLL NPs[66].

**Figure 5.**
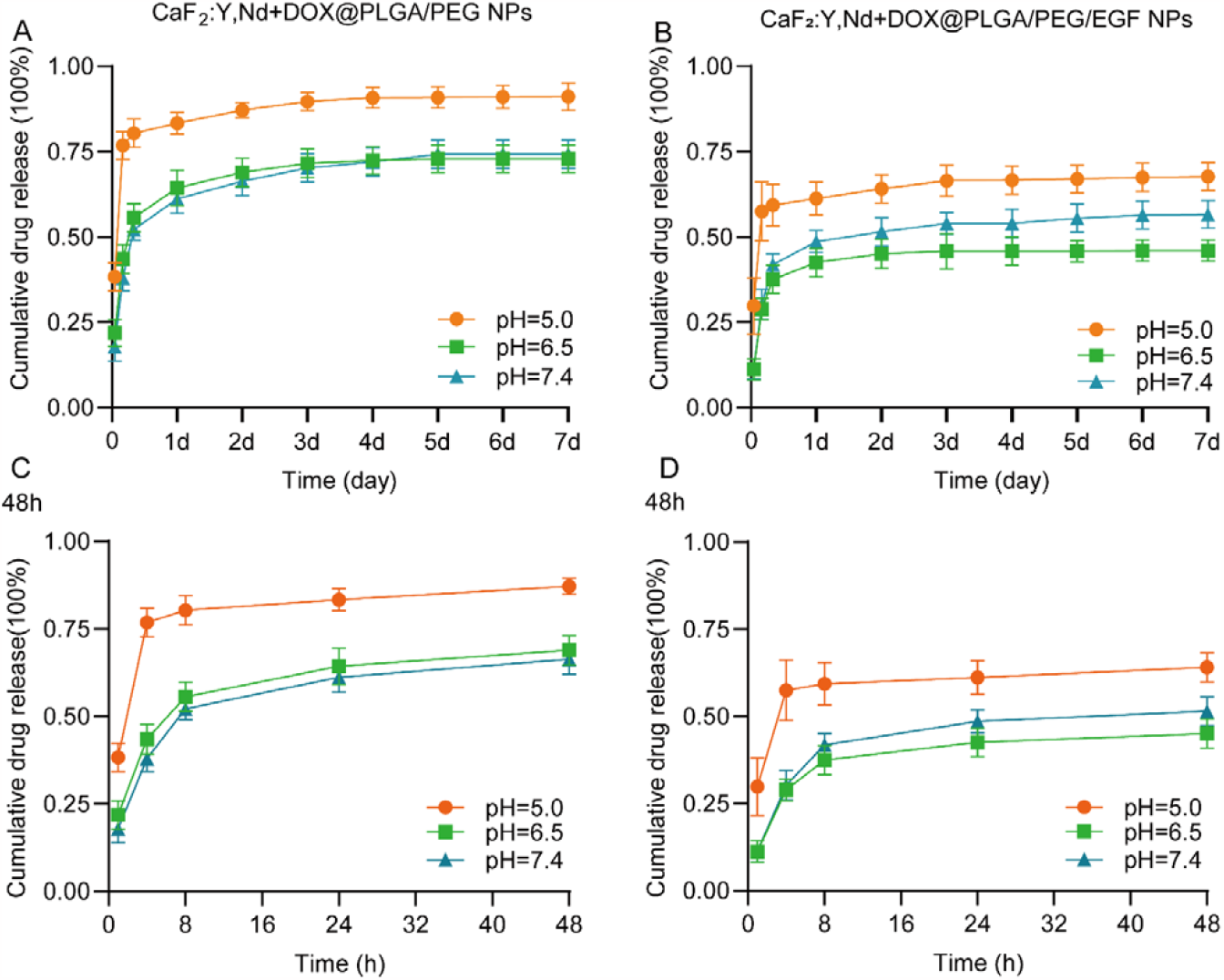
Cumulative releases of DOX from CaF_2_:Y, Nd+DOX@PLGA/PEG NPs and CaF_2_:Y, Nd+DOX@PLGA/PEG/EGF NPs in PBS at pH 5.0, pH 6.5, and pH 7.4 with 37 □ (n=3).

**Supplementary Figure 2.**
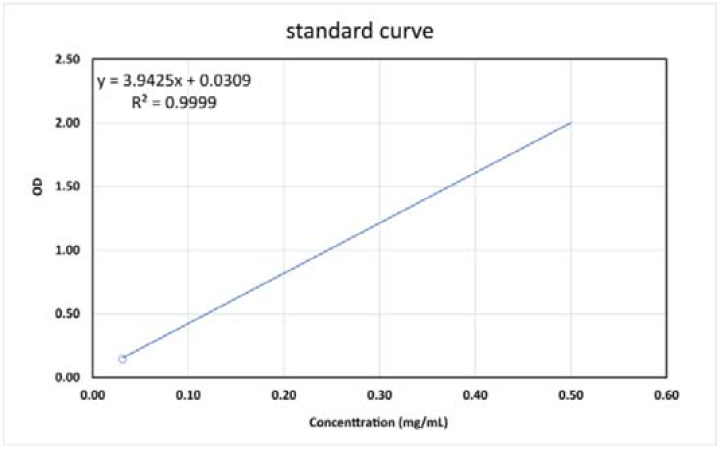
Standard curve of DOX.

**Supplementary Figure 3.**
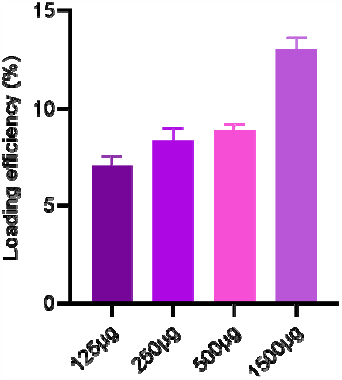
Drug loading efficiency of the drug-carriers with quantity change of DOX.

### 3.5 Cellular NP uptake

NPs are transported into the cell via the endocytic pathway, which is a time dependent process that also limits cellular uptake[67]. Effective internalization of NPs enhances their intracellular delivery and increases therapeutic effects[68]. In this study, we used the fluorescent signal of DOX to quantify the NP uptake and intracellular location by fluorescence microscopy (Figure 6A). The 4T1 cells exhibited strong punctate fluorescence, which indicated that NPs were taken up by endocytosis. As the incubation time increased, NP uptake and DOX release proportionally increased. After 48 h of incubation, we observed a numerous red fluorescent dots around the nuclei (the fluorescent dots are NPs) and a significant amount of red fluorescence overlapping with the DAPI-stained (blue) nuclei. DOX is known to promote cell death by targeting the nucleus and inhibiting topoisomerase activity[51,69]. Next, we measured the NP uptake by flow cytometry and obtained Similar to the results obtained by fluorescence microscopy, quantification of the fluorescence intensity of CaF_2_:Y Nd+DOX @ PLGA/PEG/EGF NPs in 4T1 cells by FACS analysis (Figure 6C) indicated higher cellular uptake rates of NPs with increasing incubation time.

**Figure 6.**
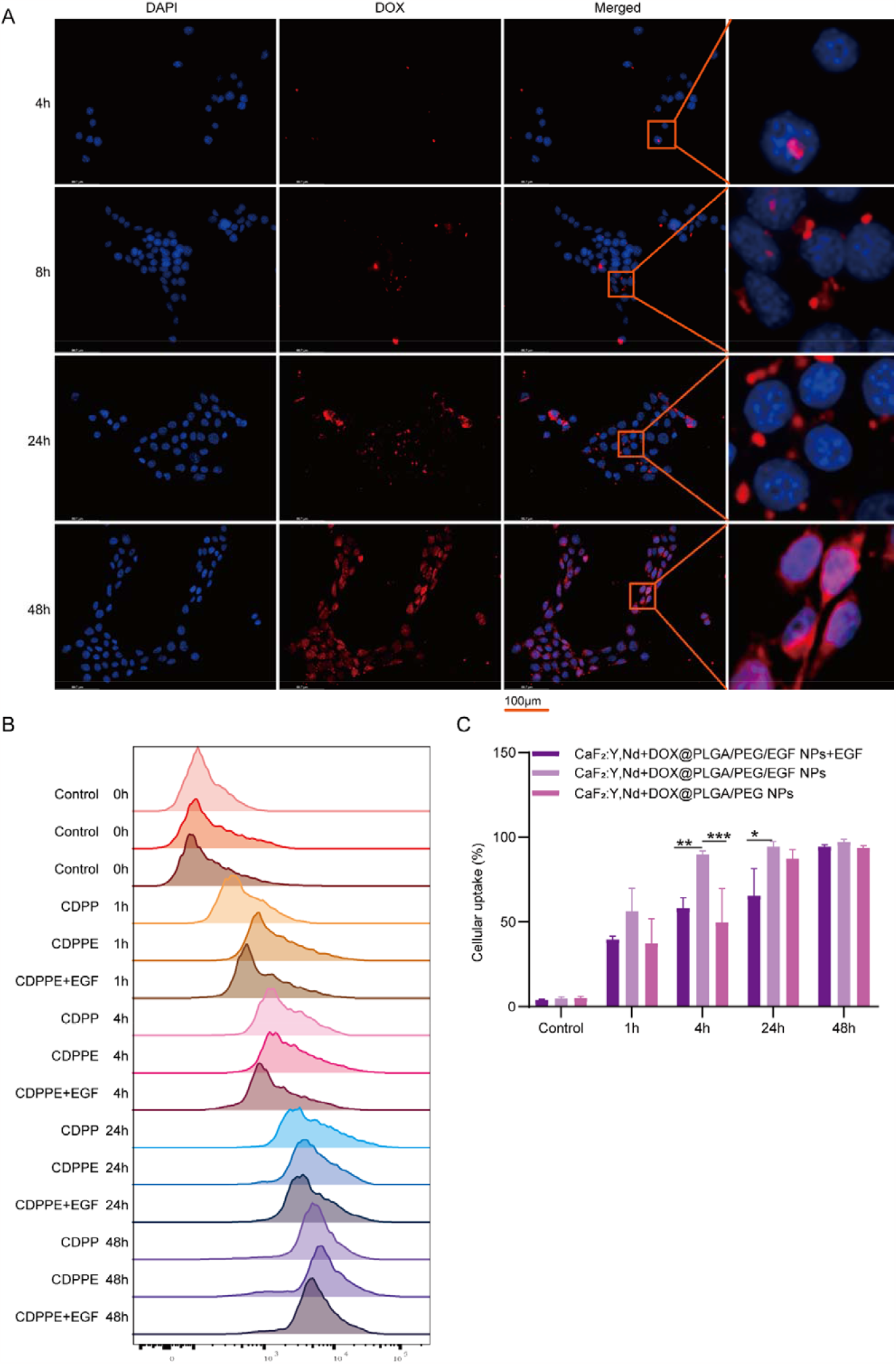
Cellular uptake studies by flow cytometry. 4T1 cells were treated with 50μg/mL CaF_2_:Y, Nd+DOX@PLGA/PEG/EGF NPs in different time points (2h, 4h, 8h, 12h, 24h and 48h). Blue (DAPI) represents nuclei; red is the DOX fluorescence. Scale bar 100 μm (A). Relative fluorescent intensity was measured during the time (C). The right shift of the chromatogram directs increased fluorescent intensity which indicates increased cellular uptake. Mean ± SD, each experiment was run triplicate. \**p*<0.05, \*\**p*<0.005,\*\*\**p*<0.0005.

EGF receptors are known to be present in large numbers on the surface of tumor cells. To determine the effect of EGF-containing NPs on cellular uptake, we treated ccells with both CaF_2_:Y Nd+DOX @ PLGA/PEG/EGF NPs and CaF_2_:Y Nd+DOX @ PLGA/PEG NPs. Flow cytometric analysis of the DOX fluorescent signal showed that after 4 h of treatment, uptake of CaF_2_:Y Nd+DOX @ PLGA/PEG/EGF NPs was significantly higher in 4T1 cancer cells, compared to CaF_2_:Y Nd+DOX @ PLGA/PEG NPs (Figure 6C). To determine the contribution of EGF to the uptake of CaF_2_:Y Nd+DOX@PLGA/PEG/EGF NPs in 4T1 cells, we performed a competition experiment. As shown in Figure 6C, the uptake of CaF_2_:Y Nd+DOX@PLGA/PEG/EGF NPs by cells cultured in medium supplemented with soluble EGF was significantly lower than of cells cultured with control medium. However, we did not observe any difference in uptake at later time points, suggesting that the presence of EGF improves the initial NP uptake.

### 3.6. *In vitro cell viabi*lity assay

To assess the cytotoxicity of the different NP formulations *in vitro, we treate*d 4T1 cells with increasing concentrations of NPs and measured cell viability using an MTS assay. The cell viability of each group was compared to the untreated control (100 % viability; Figure 7 and Supplementary Figure 4).

**Figure 7.**
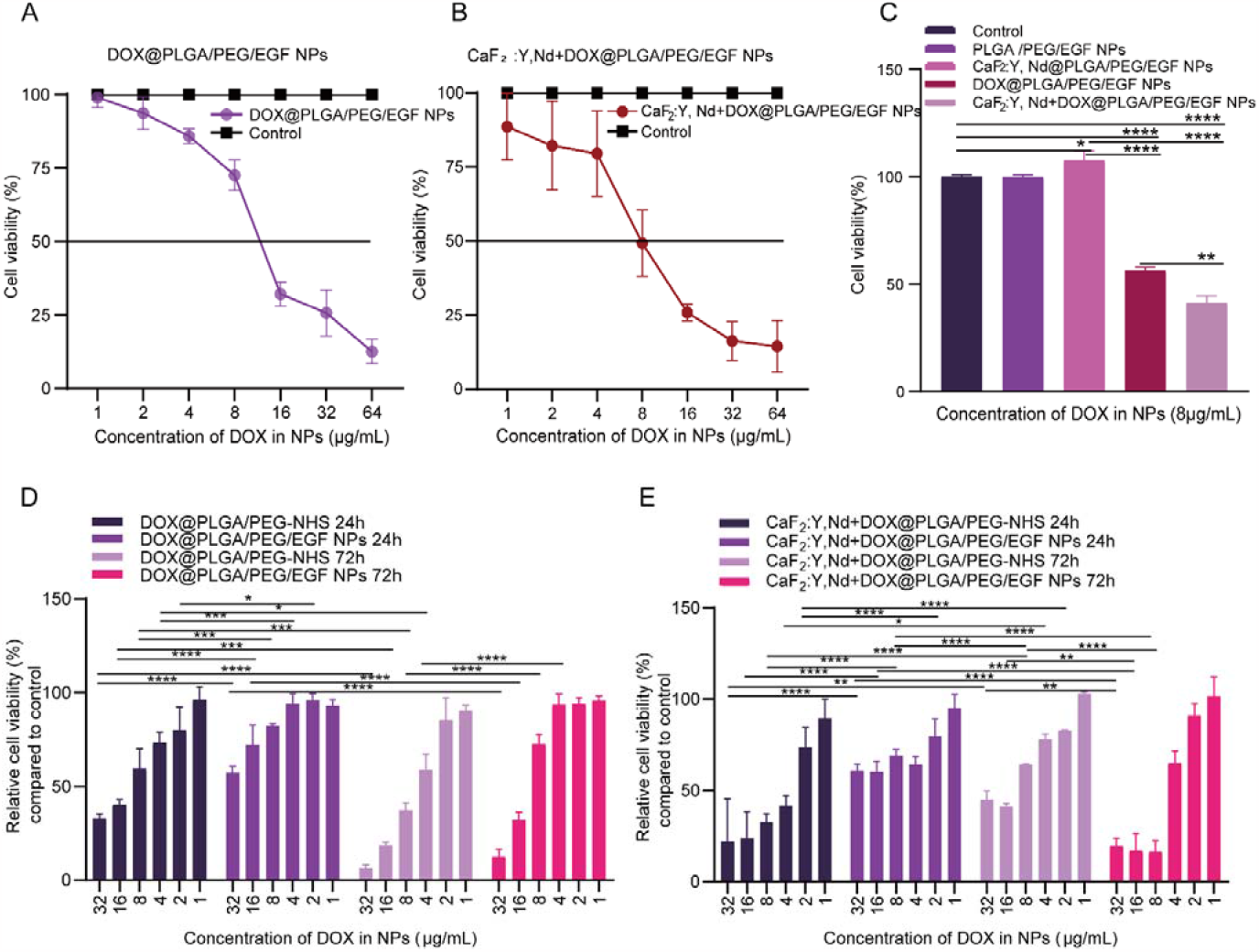
Effect of nanoparticles on cell viability. The effect on cell viability was determined by MTS after incubation with different concentrations of DOX@PLGA/PEG/EGF NPs (A) and CaF_2_:Y Nd+DOX@PLGA/PEG/EGF NPs (B). The in vitro cytotoxicity of experimental and control NPs on 4T1 cells (C). Relative viability of 4T1 cells incubated with different concentrations of NPs modified with EGF or not for 24 h and 72 h incubation via MTS assay (D-E; n=3). *p < 0.05, **p < 0.01, ***p < 0.001, ****p < 0.001 compared with each groups by t-test and Bonferroni’s multiple comparisons test.

First, we investigated whether CaF_2_:Y, Nd NPs and CaF_2_:Y, Nd@PLGA NPs in the absence of DOX (with and without EGF modification) had an effect on the viability of breast cancer cells. Treatment with different concentrations (31.25-2000 μg/mL) of PLGA/PEG NPs, PLGA/PEG/EGF NPs, CaF_2_:Y, Nd @PLGA/PEG NPs and CaF_2_:Y, Nd@PLGA/PEG/EGF NPs did not affect the viability of 4T1 cells after 24 h (Supplementary Figure 4A-B). After 72 h, CaF_2_:Y,Nd@PLGA/PEG/EGF NPs significantly induced cytotoxicity at concentrations greater than or equal to 500 μg/mL (Supplementary Figure 4 B), suggesting that RENPs were slightly cytotoxic compared to control PLGA NPs (Supplementary Figure 4A). Similar results were obtained when 4T1 cells were treated with DOX-containing NPs (Figure 7A). CaF_2_:Y,Nd+DOX @PLGA/PEG/EGF NPs were more efficient in inducing cell death in 50 % of 4T1 cells (IC50=8.032 μg/mL, equivalent to 90 μg/mL of NPs), compared to DOX@PLGA/PEG/EGF NPs (IC50=12.93 μg/mL, equivalent to 145 μg/mL of NPs).

Furthermore, CaF_2_:Y,Nd@PLGA/PEG/EGF NPs induced greater cytotoxicity than CaF_2_:Y,Nd@PLGA/PEG NPs, indicating that EGF increased the uptake of RENPs and the cytotoxicity at high concentrations (Supplementary Figure 4B).

To assess the impact of EGF-modification on cellular cytotoxicity, we treated 4T1 cells with NPs containing 8.032 μg/mL DOX (equivalent of 90 μg/mL of NPs). After 24 h, DOX@PLGA/PEG/EGF NPs reduced the survival rate of 4T1 cells by 45.9 % compared to the control group, and CaF_2_:Y, Nd+DOX@PLGA/PEG/EGF NPs significantly reduced the survival rate of 4T1 cells by 59.7 % (Figure 7 C). As the amount of NP was significantly lower than the ≥500 μg/mL of RENPs that can cause cytotoxicity in the absence of DOX (Supplementary Figure 4B), these results suggest that DOX was the major component causing cell death in DOX@PLGA/PEG/EGF NPs and CaF_2_:Y,Nd+DOX@PLGA/PEG/EGF NPs, and that CaF_2_:Y, Nd+DOX@PLGA/PEG/EGF NPs had a more significant effect on reducing cell activity.

Then, 4T1 cells were treated with NPs containing different concentrations (DOX concentrations equal to 1, 2, 4, 8, 16, and 32 μg/mL) of DOX@PLGA/PEG NPs, DOX@PLGA/PEG/EGF NPs, CaF_2_:Y, Nd+DOX@PLGA/PEG NPs and CaF_2_:Y, Nd+DOX@PLGA/PEG/EGF NPs. As shown in Figure 7D-E, after 24 h of incubation, DOX@PLGA/PEG/EGF NPs and CaF_2_:Y, Nd+DOX@PLGA/PEG/EGF NPs were less toxic than their non-EFG-modified counterparts in 4T1 cells. However, hen the incubation time was extended to 72 h, EGF-modified NPs significantly reduced the cellular activity of 4T1 cells (Figure 7 D-E). In fact, Figure 5 showed that more DOX was released more rapidly from CaF_2_:Y, Nd+DOX@PLGA/PEG NPs within the first 24 h compared to CaF_2_:Y, Nd+DOX@PLGA/PEG/EFG NPs. Thus, CaF_2_:Y, Nd+DOX@PLGA/PEG NPs were likely more effective in inducing cytotoxicity in the first 24 h due to the faster DOX release. With increasing treatment time, an increased initial cellular NP uptake due to EGF counteracted the faster release of DOX from CaF_2_:Y, Nd+DOX@PLGA/PEG NPs.

**Supplementary Figure 4.**
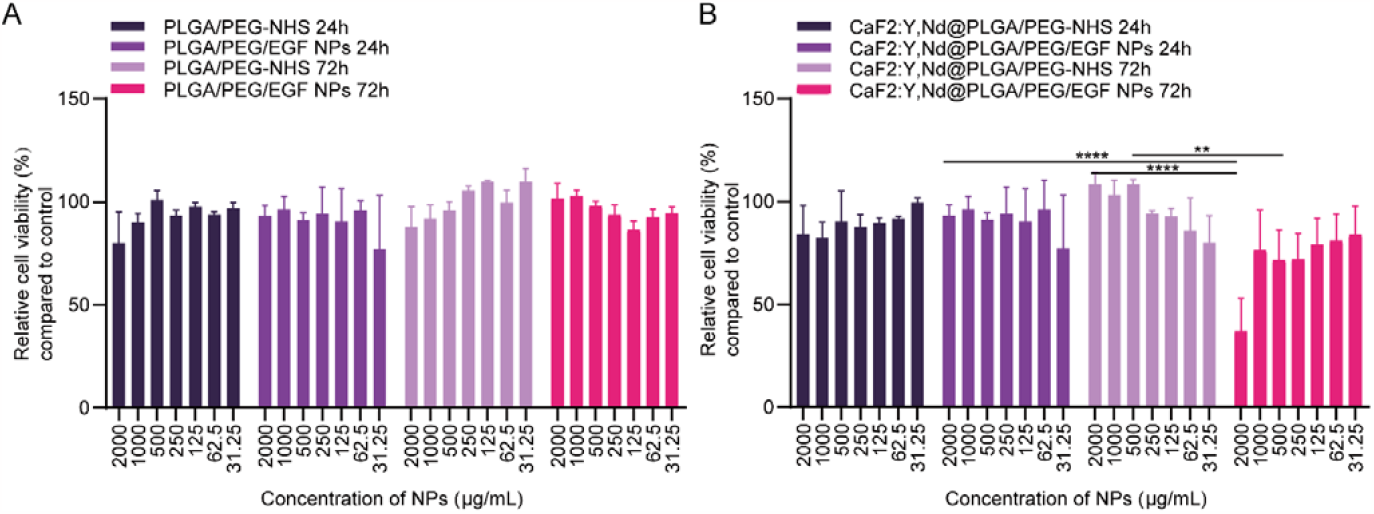
Effect of nanoparticles on cell viability. The effect on cell viability was determined by MTS after incubation with different concentrations of PLGA/PEG NPs and PLGA/ PEG/EGF NPs (A), CaF_2_:Y Nd @ PLGA/PEG NPs and CaF_2_:Y Nd@PLGA/PEG/EGF NPs (B) for 24 h and 72 h, respectively. Bonferroni’s multiple comparisons test (n=3; ** *p* < 0.01, **** *p* < 0.001.

### 3.7 NIR-II Fluorescence Imaging of NP

NIR-II fluorescence bioimaging has attracted a significant amount of attention due to its advantages over conventional imaging, such as high biosafety, high spatial resolution, strong penetration and high sensitivity. However, the effectiveness of current nanoprobes for NIR-II fluorescence imaging is poor. In this study, we explored the effectiveness of CaF_2_:Y, Nd NPs as NIR-II imaging probes. Firstly, the aqueous solution of each sample was imaged under white light. As shown in Figure 8A, CaF_2_:Y, Nd NPs appeared as white powders, which were dissolved in water and appeared as white emulsions. The powdered DOX-loaded NPs were of orange-red color, and after dissolving in water, they formed an orange-red emulsion. These results demonstrated that the NPs were water-soluble. However, we observed that CaF_2_:Y, Nd+DOX @PLGA/PEG NPs had better water solubility than CaF_2_:Y, Nd+DOX @PLGA/PEG/EGF NPs.

**Figure 8.**
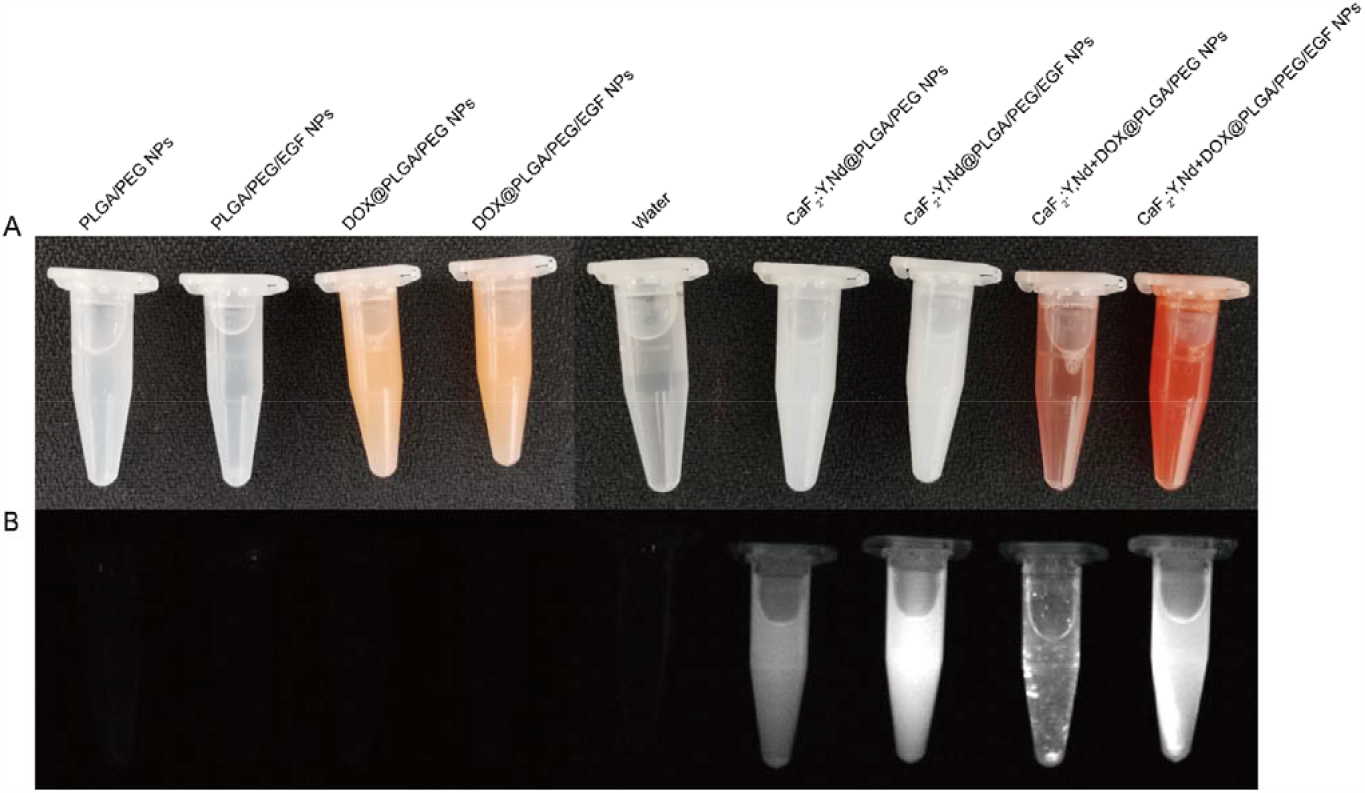
NIR-II fluorescence imaging of NPs. (A) This is the state of the aqueous solution of NPs under daylight irradiation in the experimental and control groups. (B) The representative images of NIR-II fluorescence under 808 nm laser excitation.

To verify the validity of CaF_2_:Y, Nd NPs as NIR-II imaging probes, we performed NIR-II fluorescence imaging of aqueous solutions of NPs at a concentration of 2 mg/mL using the KIS NIR-II optical imaging system (Kaer Labs, Nantes, France). As shown in Figure 8, we did not observe fluorescence for water, PLGA/PEG NPs, PLGA/PEG/EGF NPs, DOX@PLGA/PEG NPs, and DOX@PLGA/PEG/EGF NPs under excitation at 808 nm (Figure 8B). However, CaF_2_:Y, Nd@PLGA/PEG NPs, CaF_2_:Y, Nd@PLGA/PEG/EGF NPs, CaF_2_:Y, Nd+DOX @PLGA/PEG NPs, and CaF_2_:Y, Nd+DOX @PLGA/PEG/EGF NPs exhibited NIR-II fluorescence under an excitation wavelength of 808 nm (Figure 8B). This demonstrates the strong absorption of CaF_2_:Y, Nd NPs in the NIR-II window. Comparing the fluorescence intensities of CaF_2_:Y, Nd@PLGA/PEG NPs, CaF_2_:Y, Nd@PLGA/PEG/EGF NPs, CaF_2_:Y, Nd+DOX @PLGA/PEG NPs, and CaF_2_:Y, Nd+DOX @PLGA/PEG/EGF NPs, we found that CaF_2_:Y, Nd+DOX @PLGA/PEG NPs were less water-soluble, and precipitation was observed. Our data suggest that NPs carrying EGF may be more soluble in water.

## 4. Conclusion

In summary, we developed a multifunctional nanoplatform that combines chemotherapeutic drugs (DOX) with lanthanide NPs. Characterization of the NPs using TEM, DLS and UV spectrophotometry showed consistent size, shape and spectra, indicating successful NP synthesis. Cytotoxicity experiments demonstrated that encapsulation of DOX in the NPs was responsible for the cytotoxic effects in cancer cells at a NP concentration below 500 μg/mL. In addition, CaF_2_:Y Nd+DOX@PLGA/PEG/EGF NPs exhibited distinct emission peaks and a strong fluorescence response under 808 nm laser excitation, demonstrating their potential for NIR-II fluorescence imaging. Our versatile nanoplatform effectively combines the benefits of PLGA and CaF_2_:Y,Nd NPs, which not only reduces the side effects caused by DOX but also allows real-time tracking of the NP location. This also provides evidence for exploring the value of NIR-II fluorescence imaging as a probe to track the distribution of NPs in cancer therapy. In conclusion, multifunctional CaF_2_:Y Nd+DOX@PLGA/PEG/EGF NPs represent an efficient and targeted anti-cancer drug delivery system. Our drug delivery system can deliver chemotherapeutic agents and target tumor cells, as well as possesses NIR-II fluorescence imaging capabilities, which holds great promise for future applications in optical imaging detection and therapy in the medical field.

## Supplementary Information

The online version contains supplementary material available at

## Author Contributions

^#^ Contributed equally. Z.Y. and Y.H. conceived and designed the experiments. Y.H. and Z.Yu performed the experiments and responsible for the statistical analysis. Y.H. wrote the paper. T.S. performed the TEM analysis. Z.Y.,T.S., C.E. and L.J.C. revised the text. All authors have read and agreed to the published version of the manuscript.

## Acknowledgments

The authors are grateful to Kaer Labs for the valuable contributions with NIR-II imaging. The authors also thank to Roman Koning and Aat Mulder from the Koster lab at LUMC with the valuable support with electron microscopy. We would also want to express our gratitude to L. Mezzanotte from Radiology & Nuclear Medicine at Erasmus MC and Kaer Labs for their help with the NIR-II imaging.

## Informed Consent Statement

Not applicable.

## Data Availability Statement

The original contributions presented in the study are included in the article/supplementary material. Further inquiries can be directed to the corresponding author.

## References

1. Kimberly Miller, L.N., Rebecca Siegel, Ahmedin Jemal, DVM. Cancer Treatment & Survivorship Facts & Figures 2019-2021. American Cancer Society 2019.

2. Pucci, C.; Martinelli, C.; Ciofani, G. Innovative approaches for cancer treatment: current perspectives and new challenges. Ecancermedicalscience 2019, 13, 961, doi:10.3332/ecancer.2019.961.

3. Yang, P.; Quan, Z.; Hou, Z.; Li, C.; Kang, X.; Cheng, Z.; Lin, J. A magnetic, luminescent and mesoporous core-shell structured composite material as drug carrier. Biomaterials 2009, 30, 4786–4795, doi:10.1016/j.biomaterials.2009.05.038.

4. Yu, Z.; Eich, C.; Cruz, L.J. Recent Advances in Rare-Earth-Doped Nanoparticles for NIR-II Imaging and Cancer Theranostics. Front Chem 2020, 8, 496, doi:10.3389/fchem.2020.00496.

5. Patra, J.K.; Das, G.; Fraceto, L.F.; Campos, E.V.R.; Rodriguez-Torres, M.D.P.; Acosta-Torres, L.S.; Diaz-Torres, L.A.; Grillo, R.; Swamy, M.K.; Sharma, S., et al. Nano based drug delivery systems: recent developments and future prospects. J Nanobiotechnology 2018, 16, 71, doi:10.1186/s12951-018-0392-8.

6. Ruijie D. Teo†, J.T., ‡, and Harry B. Gray*,†. Lanthanides: Applications in Cancer Diagnosis and Therapy. J Med Chem 2016, 59.

7. Yu, Z.; Vepris, O.; Eich, C.; Feng, Y.; Que, I.; Camps, M.G.M.; Zhang, H.; Ossendorp, F.A.; Cruz, L.J. Upconversion nanoparticle platform for efficient dendritic cell antigen delivery and simultaneous tracking. Mikrochim Acta 2022, 189, 368, doi:10.1007/s00604-022-05441-z.

8. Kuang, Y.; Zhang, Y.; Zhao, Y.; Cao, Y.; Zhang, Y.; Chong, Y.; Pei, R. Dual-stimuli-responsive multifunctional Gd2Hf2O7 nanoparticles for MRI-guided combined chemo-/photothermal-/radiotherapy of resistant tumors. ACS Applied Materials & Interfaces 2020, 12, 35928–35939.

9. Shapoval, O.; Oleksa, V.; Slouf, M.; Lobaz, V.; Trhlikova, O.; Filipova, M.; Janouskova, O.; Engstova, H.; Pankrac, J.; Modry, A., et al. Colloidally Stable P(DMA-AGME)-Ale-Coated Gd(Tb)F3:Tb(3+)(Gd(3+)),Yb(3+),Nd(3+) Nanoparticles as a Multimodal Contrast Agent for Down- and Upconversion Luminescence, Magnetic Resonance Imaging, and Computed Tomography. Nanomaterials (Basel) 2021, 11, pdoi:10.3390/nano11010230.

10. Yu, Z.; Fu, X.; Zheng, S.; Zhang, H. Nd3+ doped LuOF nanophosphors for bimodality imaging of NIR-to-NIR-II luminescence and X-Ray computed tomography. Journal of Luminescence 2021, 231, 117753.

11. Fan, Q.; Cui, X.; Guo, H.; Xu, Y.; Zhang, G.; Peng, B. Application of rare earth-doped nanoparticles in biological imaging and tumor treatment. Journal of Biomaterials Applications 2020, 35, 237–263.

12. Zhi, G.; Song, J.; Mei, B.; Zhou, W. Synthesis and characterization of Er3+ doped CaF2 nanoparticles. Journal of alloys and compounds 2011, 509, 9133–9137.

13. Pandurangappa, C.; Lakshminarasappa, B. Morphology and optical properties of Mg and Sr doped CaF2 nanocrystals. Optics Communications 2012, 285, 2739–2742.

14. Yu, Z.; He, Y.; Schomann, T.; Wu, K.; Hao, Y.; Suidgeest, E.; Zhang, H.; Eich, C.; Cruz, L.J. Achieving Effective Multimodal Imaging with Rare-Earth Ion-Doped CaF2 Nanoparticles. Pharmaceutics 2022, 14, doi:10.3390/pharmaceutics14040840.

15. Chang, W.T.; Shih, J.Y.; Lin, Y.W.; Chen, Z.C.; Kan, W.C.; Lin, T.H.; Hong, C.S. Dapagliflozin protects against doxorubicin-induced cardiotoxicity by restoring STAT3. Arch Toxicol 2022, 10.1007/s00204-022-03298-y, doi:10.1007/s00204-022-03298-y.

16. Fu, H.Y.; Sanada, S.; Matsuzaki, T.; Liao, Y.; Okuda, K.; Yamato, M.; Tsuchida, S.; Araki, R.; Asano, Y.; Asanuma, H., et al. Chemical Endoplasmic Reticulum Chaperone Alleviates Doxorubicin-Induced Cardiac Dysfunction. Circ Res 2016, 118, 798–809, doi:10.1161/CIRCRESAHA.115.307604.

17. Siddharth, S.; Nayak, A.; Nayak, D.; Bindhani, B.K.; Kundu, C.N. Chitosan-Dextran sulfate coated doxorubicin loaded PLGA-PVA-nanoparticles caused apoptosis in doxorubicin resistance breast cancer cells through induction of DNA damage. Sci Rep 2017, 7, 2143, doi:10.1038/s41598-017-02134-z.

18. Chai, F.; Sun, L.; He, X.; Li, J.; Liu, Y.; Xiong, F.; Ge, L.; Webster, T.J.; Zheng, C. Doxorubicin-loaded poly (lactic-co-glycolic acid) nanoparticles coated with chitosan/alginate by layer by layer technology for antitumor applications. Int J Nanomedicine 2017, 12, 1791–1802, doi:10.2147/IJN.S130404.

19. Yu, X.; Sun, L.; Tan, L.; Wang, M.; Ren, X.; Pi, J.; Jiang, M.; Li, N. Preparation and characterization of PLGA–PEG–PLGA nanoparticles containing salidroside and tamoxifen for breast cancer therapy. AAPS PharmSciTech 2020, 21, 1–11.

20. Acharya, S.; Dilnawaz, F.; Sahoo, S.K. Targeted epidermal growth factor receptor nanoparticle bioconjugates for breast cancer therapy. Biomaterials 2009, 30, 5737–5750.

21. Bi, C.; Wang, A.; Chu, Y.; Liu, S.; Mu, H.; Liu, W.; Wu, Z.; Sun, K.; Li, Y. Intranasal delivery of rotigotine to the brain with lactoferrin-modified PEG-PLGA nanoparticles for Parkinson’s disease treatment. International journal of nanomedicine 2016, 11, 6547.

22. Parveen, S.; Sahoo, S.K. Polymeric nanoparticles for cancer therapy. J Drug Target 2008, 16, 108–123, doi:10.1080/10611860701794353.

23. Xu, S.; Olenyuk, B.Z.; Okamoto, C.T.; Hamm-Alvarez, S.F. Targeting receptor-mediated endocytotic pathways with nanoparticles: rationale and advances. Adv Drug Deliv Rev 2013, 65, 121–138, doi:10.1016/j.addr.2012.09.041.

24. Gocheva, G.; Ivanova, A. A Look at Receptor–Ligand Pairs for Active-Targeting Drug Delivery from Crystallographic and Molecular Dynamics Perspectives. Molecular Pharmaceutics 2019, 16, 3293–3321.

25. Stivarou, T.; Patsavoudi, E. Extracellular molecules involved in cancer cell invasion. Cancers 2015, 7, 238–265.

26. Krasinskas, A.M. EGFR Signaling in Colorectal Carcinoma. Patholog Res Int 2011, 2011, 932932, doi:10.4061/2011/932932.

27. Akbarzadeh Khiavi, M.; Safary, A.; Barar, J.; Ajoolabady, A.; Somi, M.H.; Omidi, Y. Multifunctional nanomedicines for targeting epidermal growth factor receptor in colorectal cancer. Cell Mol Life Sci 2020, 77, 997–1019, doi:10.1007/s00018-019-03305-z.

28. Yu, Z.F.; Shi, J.P.; Li, J.L.; Li, P.H.; Zhang, H.W. Luminescence enhancement of CaF2:Nd(3+) nanoparticles in the second near-infrared window for in vivo imaging through Y(3+) doping. J Mater Chem B 2018, 6, 1238–1243, doi:10.1039/c7tb03052e.

29. Pedroni, M.; Piccinelli, F.; Passuello, T.; Polizzi, S.; Ueda, J.; Haro-González, P.; Martinez Maestro, L.; Jaque, D.; García-Solé, J.; Bettinelli, M., et al. Water (H2O and D2O) Dispersible NIR-to-NIR Upconverting Yb3+/Tm3+ Doped MF2 (M = Ca, Sr) Colloids: Influence of the Host Crystal. Crystal Growth & Design 2013, 13, 4906–4913, doi:10.1021/cg401077v.

30. Bogdan, N.; Vetrone, F.; Ozin, G.A.; Capobianco, J.A. Synthesis of ligand-free colloidally stable water dispersible brightly luminescent lanthanide-doped upconverting nanoparticles. Nano Lett 2011, 11, 835–840, doi:10.1021/nl1041929.

31. Ortega-Oller, I.; Padial-Molina, M.; Galindo-Moreno, P.; O’Valle, F.; Jodar-Reyes, A.B.; Peula-Garcia, J.M. Bone Regeneration from PLGA Micro-Nanoparticles. Biomed Res Int 2015, 2015, 415289, doi:10.1155/2015/415289.

32. Wang, J.; Shi, A.; Agyei, D.; Wang, Q. Formulation of water-in-oil-in-water (W/O/W) emulsions containing trans-resveratrol. RSC Advances 2017, 7, 35917–35927, doi:10.1039/c7ra05945k.

33. Wang, L.; Sun, X. Mesoporous Silica Hybridized With Gadolinium (III) Nanoplatform for Targeted Magnetic Imaging–Guided Photothermal Breast Cancer Therapy.Dose-Response 2020, 18, 1559325820902314.

34. Ma, R.; Alifu, N.; Du, Z.; Chen, S.; Heng, Y.; Wang, J.; Zhu, L.; Ma, C.; Zhang, X. Indocyanine green-based theranostic nanoplatform for NIR fluorescence image-guided chemo/photothermal therapy of cervical cancer. International Journal of Nanomedicine 2021, 16, 4847.

35. He, Y.; Wang, M.; Fu, M.; Yuan, X.; Luo, Y.; Qiao, B.; Cao, J.; Wang, Z.; Hao, L.; Yuan, G. Iron (II) phthalocyanine loaded and AS1411 aptamer targeting nanoparticles: a nanocomplex for dual modal imaging and photothermal therapy of breast cancer. International Journal of Nanomedicine 2020, 15, 5927.

36. Gu, W.; Zhang, T.; Gao, J.; Wang, Y.; Li, D.; Zhao, Z.; Jiang, B.; Dong, Z.; Liu, H. Albumin-bioinspired iridium oxide nanoplatform with high photothermal conversion efficiency for synergistic chemo-photothermal of osteosarcoma. Drug Delivery 2019, 26, 918–927.

37. Choi, Y.; Yoon, H.Y.; Kim, J.; Yang, S.; Lee, J.; Choi, J.W.; Moon, Y.; Kim, J.; Lim, S.; Shim, M.K., et al. Doxorubicin-Loaded PLGA Nanoparticles for Cancer Therapy: Molecular Weight Effect of PLGA in Doxorubicin Release for Controlling Immunogenic Cell Death. Pharmaceutics 2020, 12, pdoi:10.3390/pharmaceutics12121165.

38. Dong, N.; Zhu, C.; Jiang, J.; Huang, D.; Li, X.; Quan, G.; Liu, Y.; Tan, W.; Pan, X.; Wu, C. Development of composite PLGA microspheres containing exenatide-encapsulated lecithin nanoparticles for sustained drug release. Asian J Pharm Sci 2020, 15, 347–355, doi:10.1016/j.ajps.2019.01.002.

39. Scott Custo1, B.B., Alex Felice1,2,3 Elisa Seria1,2. A comparative profile of total protein and six angiogenically-active growth factors in three platelet products. GMS Interdisciplinary Plastic and Reconstructive Surgery 2022.

40. Yu, Z.-f.; Shi, J.-p.; Li, J.-l.; Li, P.-h.; Zhang, H.-w. Luminescence enhancement of CaF 2: Nd 3+ nanoparticles in the second near-infrared window for in vivo imaging through Y 3+ doping. Journal of Materials Chemistry B 2018, 6, 1238–1243.

41. Bezerra, C.d.S.; Valerio, M.E. Structural and optical study of CaF2 nanoparticles produced by a microwave-assisted hydrothermal method. Physica B: Condensed Matter 2016, 501, 106–112.

42. Miller, F.; Wintzheimer, S.; Reuter, T.; Groppe, P.; Prieschl, J.; Retter, M.; Mandel, K. Luminescent Supraparticles Based on CaF2–Nanoparticle Building Blocks as Code Objects with Unique IDs. ACS Applied Nano Materials 2019, 3, 734–741.

43. Chen, Z.; Guo, N.; Ji, L.; Xu, C. Synthesis of CaF2 nanoparticles coated by SiO2 for improved Al2O3/TiC self-lubricating ceramic composites. nanomaterials 2019, 9, 1522.

44. Patel, V.K.; Saurav, J.R.; Gangopadhyay, K.; Gangopadhyay, S.; Bhattacharya, S. Combustion characterization and modeling of novel nanoenergetic composites of Co 3 O 4/nAl. RSC Advances 2015, 5, 21471–21479.

45. Huang, X.; Chen, Z.; Gao, T.; Huang, Q.; Niu, F.; Qin, L.; Huang, Y. Hydrogen Generation by Hydrolysis of an Al/Al2O3-Composite Powder After Heat Treatment. Energy Technology 2013, 1, 751–756.

46. Nidhin, M.; Sreeram, K.J.; Nair, B.U. Green synthesis of rock salt CoO nanoparticles for coating applications by complexation and surface passivation with starch. Chemical Engineering Journal 2012, 185, 352–357.

47. Muhammad, W.; Ullah, N.; Haroon, M.; Abbasi, B.H. Optical, morphological and biological analysis of zinc oxide nanoparticles (ZnO NPs) using Papaver somniferum L. RSC advances 2019, 9, 29541–29548.

48. Sisubalan, N.; Ramkumar, V.S.; Pugazhendhi, A.; Karthikeyan, C.; Indira, K.; Gopinath, K.; Hameed, A.S.H.; Basha, M.H.G. ROS-mediated cytotoxic activity of ZnO and CeO2 nanoparticles synthesized using the Rubia cordifolia L. leaf extract on MG-63 human osteosarcoma cell lines. Environmental Science and Pollution Research 2018, 25, 10482–10492.

49. Zhao, J.; Yang, H.; Li, J.; Wang, Y.; Wang, X. Fabrication of pH-responsive PLGA(UCNPs/DOX) nanocapsules with upconversion luminescence for drug delivery. Sci Rep 2017, 7, 18014, doi:10.1038/s41598-017-16948-4.

50. Lei, T.; Srinivasan, S.; Tang, Y.; Manchanda, R.; Nagesetti, A.; Fernandez-Fernandez, A.; McGoron, A.J. Comparing cellular uptake and cytotoxicity of targeted drug carriers in cancer cell lines with different drug resistance mechanisms. Nanomedicine 2011, 7, 324–332, doi:10.1016/j.nano.2010.11.004.

51. Hueso, D.; Fontecha, J.; Gomez-Cortes, P. Comparative study of the most commonly used methods for total protein determination in milk of different species and their ultrafiltration products. Front Nutr 2022, 9, 925565, doi:10.3389/fnut.2022.925565.

52. Kang, X.; Yang, D.; Dai, Y.; Shang, M.; Cheng, Z.; Zhang, X.; Lian, H.; Lin, J. Poly (acrylic acid) modified lanthanide-doped GdVO 4 hollow spheres for up-conversion cell imaging, MRI and pH-dependent drug release. Nanoscale 2013, 5, 253–261.

53. Li, X.; Shen, D.; Yang, J.; Yao, C.; Che, R.; Zhang, F.; Zhao, D. Successive layer-by-layer strategy for multi-shell epitaxial growth: shell thickness and doping position dependence in upconverting optical properties. Chemistry of Materials 2013, 25, 106–112.

54. Abel, K.A.; Boyer, J.-C.; Veggel, F.C.v. Hard proof of the NaYF4/NaGdF4 nanocrystal core/shell structure. Journal of the American Chemical Society 2009, 131, 14644–14645.

55. Su, Q.; Han, S.; Xie, X.; Zhu, H.; Chen, H.; Chen, C.-K.; Liu, R.-S.; Chen, X.; Wang, F.; Liu, X. The effect of surface coating on energy migration-mediated upconversion. Journal of the American Chemical Society 2012, 134, 20849–20857.

56. Zhang, F.; Che, R.; Li, X.; Yao, C.; Yang, J.; Shen, D.; Hu, P.; Li, W.; Zhao, D. Direct imaging the upconversion nanocrystal core/shell structure at the subnanometer level: shell thickness dependence in upconverting optical properties. Nano letters 2012, 12, 2852–2858.

57. Mittal, G.; Sahana, D.K.; Bhardwaj, V.; Ravi Kumar, M.N. Estradiol loaded PLGA nanoparticles for oral administration: effect of polymer molecular weight and copolymer composition on release behavior in vitro and in vivo. J Control Release 2007, 119, 77–85, doi:10.1016/j.jconrel.2007.01.016.

58. Tsai, L.H.; Yen, C.H.; Hsieh, H.Y.; Young, T.H. Doxorubicin Loaded PLGA Nanoparticle with Cationic/Anionic Polyelectrolyte Decoration: Characterization, and Its Therapeutic Potency. Polymers (Basel) 2021, 13, pdoi:10.3390/polym13050693.

59. Rawat, P.S.; Jaiswal, A.; Khurana, A.; Bhatti, J.S.; Navik, U. Doxorubicin-induced cardiotoxicity: An update on the molecular mechanism and novel therapeutic strategies for effective management. Biomed Pharmacother 2021, 139, 111708, doi:10.1016/j.biopha.2021.111708.

60. Yu, T.; Li, Y.; Gu, X.; Li, Q. Development of a Hyaluronic Acid-Based Nanocarrier Incorporating Doxorubicin and Cisplatin as a pH-Sensitive and CD44-Targeted Anti-Breast Cancer Drug Delivery System. Front Pharmacol 2020, 11, 532457, doi:10.3389/fphar.2020.532457.

61. Wang, C.; Zhao, T.; Li, Y.; Huang, G.; White, M.A.; Gao, J. Investigation of endosome and lysosome biology by ultra pH-sensitive nanoprobes. Adv Drug Deliv Rev 2017, 113, 87–96, doi:10.1016/j.addr.2016.08.014.

62. Huynh, K.K.; Grinstein, S. Regulation of vacuolar pH and its modulation by some microbial species. Microbiol Mol Biol Rev 2007, 71, 452–462, doi:10.1128/MMBR.00003-07.

63. Danhier, F.; Ansorena, E.; Silva, J.M.; Coco, R.; Le Breton, A.; Preat, V. PLGA-based nanoparticles: an overview of biomedical applications. J Control Release 2012, 161, 505–522, doi:10.1016/j.jconrel.2012.01.043.

64. Zheng, F.; Wang, S.; Shen, M.; Zhu, M.; Shi, X. Antitumor efficacy of doxorubicin-loaded electrospun nano-hydroxyapatite–poly (lactic-co-glycolic acid) composite nanofibers. Polymer Chemistry 2013, 4, 933–941.

65. Li, Y.; Leng, Q.; Pang, X.; Shi, H.; Liu, Y.; Xiao, S.; Zhao, L.; Zhou, P.; Fu, S. Therapeutic effects of EGF-modified curcumin/chitosan nano-spray on wound healing. Regen Biomater 2021, 8, rbab009, doi:10.1093/rb/rbab009.

66. Wang, Y.; Liu, P.; Qiu, L.; Sun, Y.; Zhu, M.; Gu, L.; Di, W.; Duan, Y. Toxicity and therapy of cisplatin-loaded EGF modified mPEG-PLGA-PLL nanoparticles for SKOV3 cancer in mice. Biomaterials 2013, 34, 4068–4077.

67. Thorn, C.F.; Oshiro, C.; Marsh, S.; Hernandez-Boussard, T.; McLeod, H.; Klein, T.E.; Altman, R.B. Doxorubicin pathways: pharmacodynamics and adverse effects. Pharmacogenet Genomics 2011, 21, 440–446, doi:10.1097/FPC.0b013e32833ffb56.

68. Noorani, L.; Pourgholami, M.H.; Liang, M.; Morris, D.L.; Stenzel, M. Albendazole loaded albumin nanoparticles for ovarian cancer therapy. European Journal of Nanomedicine 2014, 6, doi:10.1515/ejnm-2014-0026.

69. Zeng, Y.; Zhang, X.; Lin, D.; Feng, X.; Liu, Y.; Fang, Z.; Zhang, W.; Chen, Y.; Zhao, M.; Wu, J., et al. A lysosome-targeted dextran-doxorubicin nanodrug overcomes doxorubicin-induced chemoresistance of myeloid leukemia. J Hematol Oncol 2021, 14, 189, doi:10.1186/s13045-021-01199-8.

